# Neuronal origin of the temporal dynamics of spontaneous BOLD activity correlation

**DOI:** 10.1101/169698

**Authors:** Teppei Matsui, Tomonari Murakami, Kenichi Ohki

## Abstract

Resting-state functional connectivity (FC) has become a major fMRI method to study network organization of human brains. There has been recent interest in the temporal fluctuations of FC calculated using short time windows (“dynamic FC”) because this method could provide information inaccessible with conventional “static” FC, which is typically calculated using the entire scan lasting several tens of minutes. Although multiple studies have revealed considerable temporal fluctuations in FC, it is still unclear whether the fluctuations of FC measured in hemodynamics reflect the dynamics of underlying neural activity. We addressed this question using simultaneous imaging of neuronal calcium and hemodynamic signals in mice and found coordinated temporal dynamics of calcium FC and hemodynamic FC measured in the same short time windows. Moreover, we found that variation in transient neuronal coactivation patterns (CAPs) was significantly related to temporal fluctuations of sliding window FC in hemodynamics. Finally, we show that the observed dynamics of FC cannot be fully accounted for by simulated data assuming stationary FC. These results provide evidence for the neuronal origin of dynamic FC and further suggest that information relevant to FC is condensed in temporally sparse events that can be extracted using a small number of time points.

## Introduction

Resting-state functional connectivity (FC) uses temporal correlation of spontaneous neuronal activity to assess network organization of brain regions in a non-invasive manner (Fox and Raichle 2007). Traditionally, FC has been calculated using all time points in a scan that typically lasts between several minutes to tens of minutes (Biswal et al. 1995; Fox et al. 2005; Van Dijk et al. 2010). Such “static” FC has been shown to largely reflect anatomical connectivity (Adachi et al. 2012; Honey et al. 2009; Matsui et al. 2012; Matsui et al. 2011; Vincent et al. 2007). Recently, in contrast to the traditional analysis of “static” FC, the temporal fluctuation of FC across short time windows is increasingly recognized as a useful aspect of FC (Allen et al. 2014; Hutchison et al. 2013; Zalesky et al. 2014). Such “dynamic” FC (dFC) calculated using short time windows could provide information that is inaccessible with static FC about the functional network organizations of healthy and diseased brains (Calhoun et al. 2014; Preti et al. 2016) [We note that the term “dynamics” refers to the non-stationarity of FC obtained with the sliding window analyses and does not refer to a process that is not invariant against to temporal reordering of the samples (Liégeois et al. 2017)]. The presence of temporal fluctuations in FC has also informed theoreticians on how to constrain realistic models of brain networks (Deco et al. 2013; Hansen et al. 2015; Messé et al. 2014).

However, despite growing interest, the neurophysiological basis of dFC is still weak. Previous attempts to investigate the neural origin of dFC by simultaneous measurement of electrophysiological and functional magnetic resonance imaging (fMRI) have been limited in several ways (Lu et al. 2007; Pan et al. 2011; Tagliazucchi et al. 2012b; Thompson et al. 2013). In some studies, electrophysiological recording was limited to a small number of recording sites due to technical difficulty (Lu et al. 2007; Pan et al. 2011; Thompson et al. 2013); hence, information on the global pattern of neuronal activity was lacking. In another study, electrophysiological signals were obtained with an electroencephalogram, which records global neuronal activity but lacks precise spatial information (Tagliazucchi et al. 2012b). Thus, the link between temporal fluctuations of FC in hemodynamics and that of large-scale neuronal activity has not been adequately proven.

Several studies have also questioned whether the apparent “dynamics” of FC calculated using the sliding window method is related to temporal instability of spontaneous brain network (Hindriks et al. 2016; Laumann et al. 2016). Whereas many studies have attributed temporal fluctuations of sliding window FC to the non-stationarity of spontaneous neuronal activity correlation (Allen et al. 2014; Zalesky et al. 2014), some recent studies have demonstrated that the temporal fluctuations of FC observed in the real and the simulated data, which is stationary by construction, are statistically indistinguishable (Hindriks et al. 2016; Laumann et al. 2016). Furthermore, Laumann and colleagues have shown that, in the human resting state, blood oxygen level dependent (BOLD) time series are largely stationary, discounting head-motion and fluctuating arousal (Laumann et al. 2016). Therefore, not only the neuronal basis of dFC, but also the existence of the statistical non-stationarity of FC, or at least the capability of sliding window methods to detect the non-stationarity, is called into question.

In the present study, we addressed these questions using simultaneous imaging of neuronal calcium and BOLD hemodynamic signals in the entire neocortex of transgenic mice expressing a genetically encoded calcium indicator (Matsui et al. 2016; Vanni and Murphy 2014; White et al. 2011). In the present experimental setup, the wide-field calcium signal provided access to neuronal activity at higher temporal resolution and signal-to-noise ratio compared to that of the hemodynamic signal (Matsui et al. 2016; Murakami et al. 2017; Murakami et al. 2015; Tohmi et al. 2014; Vanni and Murphy 2014). Moreover, unlike human fMRI data, in the present dataset, mice were tightly head-fixed and lightly anesthetized; thus, excluding head motions from contaminating FC. The main findings of the present study were as follows. First, we found consistency between the “dynamics” of FC calculated using calcium and hemodynamic signals, suggesting a neuronal origin of the temporal fluctuations of hemodynamic FC. Second, we found that temporal fluctuations of the spatial pattern of transient neuronal coactivations as measured in the calcium signal were significantly correlated with temporal fluctuations of hemodynamic FC. Finally, we found that the statistical properties of sliding window FC were significantly different between the real and the simulated data suggesting non-stationarity of resting-state FC.

## Materials and Methods

### Animals

Emx1-IRES-cre and Ai38 (Zariwala et al. 2012) mice were obtained from the Jackson Laboratory (Sacramento, CA). These mice were crossed and all cortical excitatory neurons expressed GCaMP3. Mice (P60–P90) were prepared for *in vivo* wide-field simultaneous imaging. Anesthesia was induced with isoflurane (3 %) and maintained with isoflurane (1 – 2 % in surgery, 0.5 – 0.8 % during imaging) and chlorprothixene (0.3 – 0.8 mg/kg, intramuscular injection). For simultaneous imaging of calcium and hemodynamic signals, a custom-made metal head plate was attached to the skull using dental cement (Sun Medical Company, Ltd, Shiga, Japan) and a large craniotomy was made over the whole cortex. The craniotomy was sealed with 1 % agarose and a glass coverslip. During the imaging, body temperature was maintained by a heat pad. All experiments were carried out in accordance with the NIH Guide for the Care and Use of Laboratory Animals, the institutional animal welfare guidelines set forth by the Animal Care and Use Committee of Kyushu University, and the study was approved by the Ethical Committee of Kyushu University.

### Simultaneous Calcium and Intrinsic Signal Imaging

The data for simultaneous imaging of calcium and hemodynamic signals was taken from a published report (Matsui et al. 2016). Briefly, simultaneous imaging of calcium and intrinsic signals *in vivo* was performed using a macro zoom fluorescence microscope (MVX-10, Olympus, Tokyo, Japan) or an upright fluorescence microscope (ECLIPSE Ni-U, Nikon, Tokyo, Japan), equipped with a 1× objective. A 625 nm LED light source was used to obtain intrinsic signals, which we referred to as the hemodynamic signal (Vanni and Murphy 2014). At this wavelength, the optical intrinsic signal primarily reflects deoxyhemoglobin signal (HbR) (Ma et al. 2016). GCaMP was excited by a 100 W mercury lamp through a GFP mirror unit (Olympus, Tokyo, Japan). Intrinsic signal data was collected at a frame rate of 5 Hz using a CCD camera (1,000m; Adimec, Boston, MA, U.S.A.) and calcium signal data was collected at a frame rate of 10 Hz using a CCD camera (DS-Qi1 Mc; Nikon, Tokyo, Japan). The emission filters were 625 nm long pass (SC-60, Fuji film, Tokyo, Japan) for intrinsic signals, and 505-535 nm band pass (FF01-520/35-25, Semrock, Lake Forest, Illinois) for calcium signals. Data were acquired for 30-60 min per animal (5 min per scan).

### Data Preprocessing

All data analyses were conducted in Matlab (MathWorks, Natick, MA) using a method described previously (Matsui et al. 2016). Briefly, all the image frames were corrected for possible within-scan motion by rigid-body transformation. Calcium and hemodynamic images were then coregistered by rigid-body transformation using manually selected anatomical landmarks that were visible in both images (e.g., branching points of blood vessels). All of the images were then spatially down-sampled by a factor of two. Pixels within the cortex (at this point including large blood vessels including the sinus) were extracted manually. For both calcium and hemodynamics, slow drift in each pixel’s time course was removed using a high-pass filter (> 0.01 Hz, second order Butterworth. No low-pass filter was used). After filtering, each pixel’s time course was normalized by subtracting the mean across time and then dividing by the standard deviation across time. Global signal regression was conducted by regressing out the time course of average signal within the brain from each pixel’s time course. Finally, hemodynamic signal was multiplied by ‐1 to set the polarity of the activity change equal to that in the calcium signal.

In some analyses, the calcium signal was further preprocessed. To obtain the high frequency calcium signal, an additional high-pass filter (> 0.1 Hz) followed by a median filter was applied. The median filter was applied as follows: For each frame, we defined a time-window (width = 200 frames) whose center was positioned on the frame. Then, the signal of the frame was replaced by the median of the time-window [B(t) = median(A(t-100), …, A(t+100)); where A(k) denotes the original signal at frame k and B(k) denotes the median filtered signal at frame k]. To obtain the low frequency calcium signal, an additional low-pass filter (< 0.1 Hz) was applied.

### Extraction of Region-of-Interest (ROI) Time Courses

Selection of ROI and time courses are conducted as described previously (Matsui et al. 2016). Briefly, 38 cortical regions (19 for each hemisphere) were selected as ROIs based on a previous mouse functional connectivity study (White et al. 2011) (Supplementary Fig. 1). Each ROI was a 6 × 6 pixel square (0.5 mm × 0.5 mm) centered at a selected coordinate. The time course for each ROI was calculated by averaging the time courses of pixels within the ROI that corresponded to gray matter. ROIs located outside of the FOV were discarded.

### Analysis of FC

For both calcium and hemodynamic signals, FC was calculated using a standard seed-based correlation method (Matsui et al. 2016). First, the correlation coefficient between the time course of a selected ROI ("seed time course") and the time course of every pixel within the brain was calculated. Second, FC values were averaged across scans to obtain FC values for each pixel. The spatial correlation between FC maps of calcium and hemodynamic signals was calculated by taking the pixel-by-pixel correlation coefficient between the two maps using all the gray matter pixels. FC with short time window was obtained by taking correlation coefficient using all the frames within a 30 sec window. Steps of 3 sec and 30 sec were used for the sliding window and non-overlapping window, respectively. Scan-shifted control was calculated by shifting the scan number of hemodynamics data relative to simultaneously obtained calcium data.

### Analysis of Co-Activation Patterns (CAPs)

CAP analysis was adopted from previous fMRI studies (Liu and Duyn 2013). Briefly, the calcium time course from each ROI was z-normalized. CAPs were calculated for each ROIs. Frames corresponding to large peaks (> 2 s.d.) of the time course of a given ROI were considered CAPs. We examined whether CAPs calculated using the calcium signal could predict the sliding window FC calculated using the hemodynamic signal. For each ROI and each time window, we quantified the similarity between the sliding window FC and CAPs by calculating the spatial correlation between the FC map and CAPs. To quantify coordinated temporal variations in CAPs and sliding-window FC exist, we also calculated the temporal deviation of CAP and FC from the mean patterns as described in the following. First, we calculated average patterns of CAPs and FC in a given scan. For a given scan and each ROI_*i*_, the average of all CAPs was calculated using the entire scan[(CAP_*scan*_)_*i*_]. Similarly, an FC map was calculated for the same scan and the same ROI [(FC_*scan*_)_*i*_]. We repeated this procedure to obtain ROI-by-ROI matrices of CAP_*scan*_ and FC_*scan*_. Next, we calculated ROI-by-ROI matrices of CAPs and FC in a short time window (CAP_*window*_ and FC_*window*_) using the same procedure used to calculate CAP_*scan*_ and FC_*scan*_ but for each 30 sec time window. Then, the difference between CAP_*window*_ and CAP_*scan*_ was taken to quantify the deviation of CAP in a time window from the mean CAP pattern in the entire scan [ΔCAP = CAP_*window*_ – CAP_*scan*_]. Similarly, ΔFC was obtained by subtracting FC_*scan*_ from FC_*window*_. Finally, the correlation coefficient between non-diagonal elements of the ROI-by-ROI matrices of ΔCAP and ΔFC were calculated. When CAPs were absent for a particular ROI in a time window, that ROI was omitted from the calculation for the time window.

### Cluster Analysis and Kurtosis Analysis

For the state analysis of sliding window FC, we adopted the k-means clustering algorithm used in the previous studies (Allen et al. 2014; Laumann et al. 2016). Correlation distance (1-r) was used to compute the separation between each window’s FC-matrix (using all 38 ROIs) and the k-means clustering was iterated 100 times with random centroid positions to avoid local minima. The windowed FC-matrices were mean-centered by scan to eliminate scan-level and subject-level features from contributing the clustering result. The k-means clustering was applied in the same manner to the simulated data that was matched in size to the real data. The cluster validity index was used to evaluate the quality of clustering for the range of cluster numbers (k = 2–10). The cluster validity index was computed as the average ratio of within-cluster distance to between-cluster distance.

The non-stationarity of spontaneous neuronal signal correlation was assessed by calculating the multivariate kurtosis using the same procedure as described by Laumann and colleagues (Laumann et al. 2016). One value of kurtosis was calculated for each FC matrix (using all 38 ROIs) obtained at each scan. The same procedure was applied to the simulated data that was matched in size to the real data. A significant difference between the kurtosis measure of the real and the simulated data indicates either non-stationarity of FC or non-Gaussianity of the signal or both.

### Time Course Simulation

To obtain a null dataset to evaluate the non-stationarity of the real data, we constructed simulated time courses using a method developed by Laumann and colleagues (Laumann et al. 2016). Briefly, random normal deviates having the same dimensionality as a real dataset are sampled. These time courses are multiplied in the spectral domain by the average power spectrum of the (bandpass filtered) real data. These time courses are then projected onto the eigenvectors derived from the covariance matrix of the real data. This procedure produces simulated data that are stationary by construction but matched to real data in the covariance structure and mean spectral content.

### Assessment of motion-related and physiological artifacts

To estimate the strength of motion-related and physiological artifacts from functional images, we calculated DVARS using the hemodynamic signal with a procedure described in fMRI literatures (Power et al. 2012; Power et al. 2014). Specifically, for each scan, after the preprocessing, root mean squares of the temporal derivatives of the hemodynamic time courses were calculated and averaged across ROIs to obtain one time course of DVARS. To remove the data potentially contaminated by the artifacts, we conducted two types of analyses. First, for each scan, the histogram of DVARS was calculated to exclude scans with strong skewness (Supplementary Fig. 3) (scan-censoring). To define a DVARS histogram with strong skewness, we calculated DVARS histograms for 1000 sets of the simulation described in the preceding section. If the skewness of the data was larger than the 99^th^ percentiles of the simulation, the scan was considered to be strongly skewed. Second, frame-censoring was conducted at multiple DVARS thresholds as described in the fMRI literatures (Power et al. 2012; Power et al. 2014). For a given threshold, frames with DVARS larger than the threshold were simply discarded from the subsequent analyses.

### Assessment of the normality of the signal

To check for the normality of the hemodynamic and the calcium signals, we computed the 4^th^ moment of the signal distribution for each scan. For each scan, signals in all the ROIs were concatenated before calculating the 4^th^ moment. Similar results were obtained when the 4^th^ moment was calculated separately for individual ROIs and then the mean of the 4^th^ moment was obtained for each scan (data not shown). The calculated 4^th^ moment of the real data was then compared with that of the 1000 simulated time courses obtained by the procedure described above. If the 4^th^ moment of the real data was larger than the 99^th^ percentile of the simulation, the signal in the scan was considered to be non-normal. In the case of the calcium signal, we applied a median filter to remove high frequency noise and enforce the normality of the signal. Calcium time courses often contained some high frequency noise on top of the slower calcium activity (Fig. 1A), which was most likely due to the moderate level of excitation light adjusted to avoid bleaching of GCaMP fluorophore. We consider that the median filter removed this high frequency noise while retaining the slower calcium signal that was approximately normally distributed.

**Figure 1.**
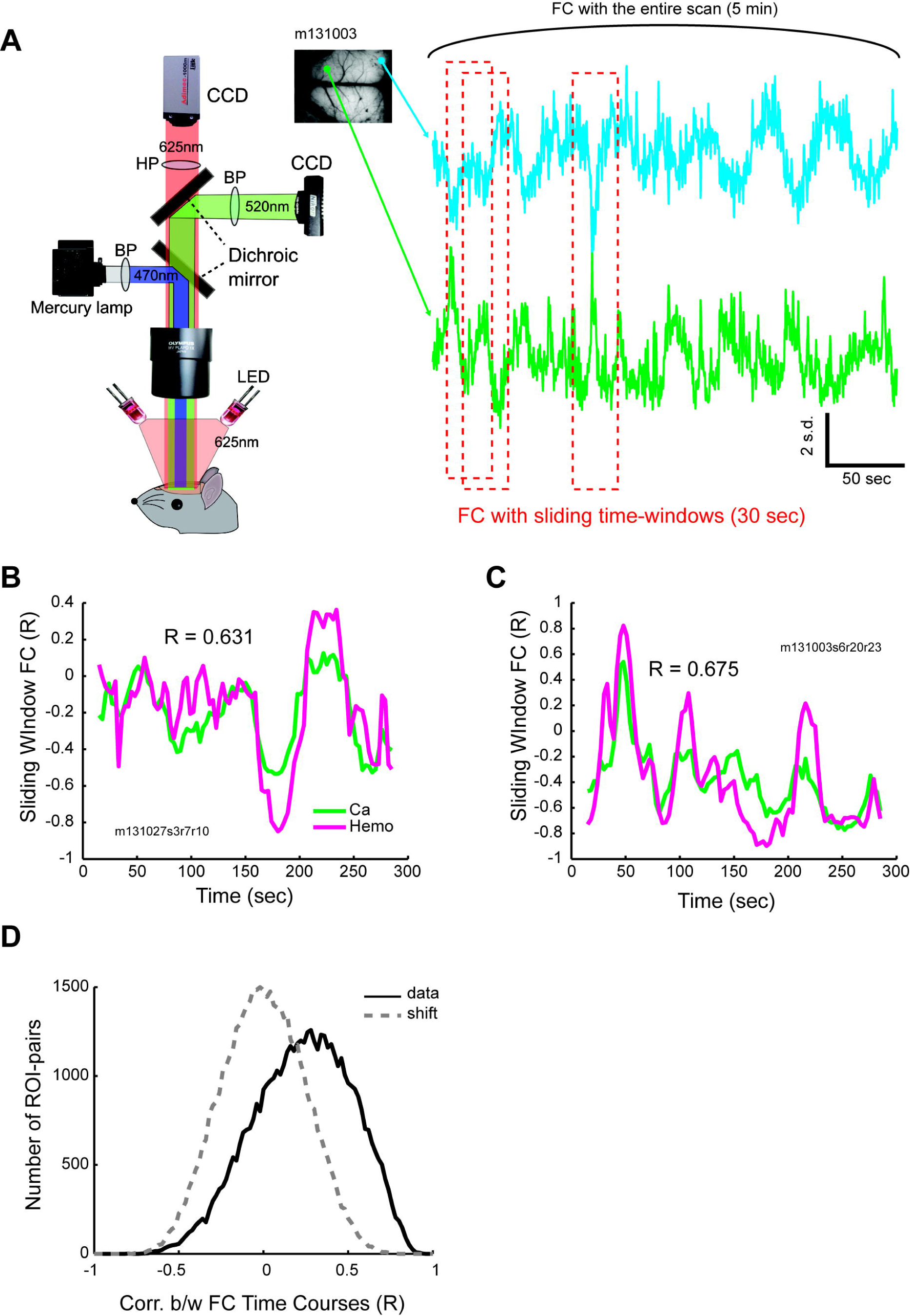
Representative dynamics of simultaneously observed calcium and hemodynamic FC. **(A)** Experimental setup. The left most panel shows the setup for simultaneous imaging. Right side shows example calcium time courses for two ROIs (green and cyan traces indicate M1 and V1 ROIs, respectively. Positions of the ROIs indicated in the example field of view. See Supplementary Figure 1 for abbreviations). FC with short time window uses subset of frames contained in short (30 sec) windows (red dotted squares). Sliding FC for hemodynamic signal was carried out similarly. **(B)-(C)** Examples dynamics of calcium and hemodynamic FC. (B) FC between right V1 and right AC. (C) FC between left M1 and left pPar. See Supplementary Figure 1 for ROI positions and abbreviations. **(D)** Histogram of correlation between time courses of Ca-FC and Hemo-FC for the data (solid line) and the scan-shifted control (dotted line). Data from all pairs of ROIs for all scans obtained in all mice were used.

## Results

### Consistent FC dynamics in calcium and hemodynamic signals

Transgenic mice expressing GCaMP in neocortical neurons were used to simultaneously measure the neuronal calcium signal and hemodynamics in a large portion of the bilateral neocortex (Fig. 1A) (Matsui et al. 2016). Mice were lightly anesthetized and head-fixed with metal plates so that head-motion could not contaminate the signals. For both calcium and hemodynamic signals, power spectra of the signals exhibited an approximately linear trend on log-log plots (Supplementary Fig. 2) suggesting that the non-neuronal artifact was small. We used sliding window correlation (30 sec window at 3 sec steps) to examine whether the calcium and hemodynamic FC in mice exhibited “dynamic” changes. Consistent with previous reports in humans (Allen et al. 2014; Chang and Glover 2010; Zalesky et al. 2014) and other animals (Hutchison et al. 2014; Majeed et al. 2009), FC between pairs of ROIs calculated with sliding windows showed considerable variability over different time points both in calcium signal and hemodynamics (Fig. 1B–C). Consistent with the idea that variability in hemodynamic FC arises from underlying neuronal activity, we found close matches between dFC of calcium and hemodynamics (correlation coefficients, 0.631 and 0.675 for Figs. 1B and 1C, respectively). Correlation between the time courses of calcium FC and hemodynamic FC was significantly larger for the data than for the scan-shifted control (P < 10^-20^, Kolmogorov-Smirnov test; Fig. 1D).

To further examine the consistency between dFC in the calcium signal and hemodynamics in the entire neocortex, we calculated the calcium and hemodynamic FC among all pairs of ROIs and compared them across time windows (Fig. 2A–B). The ROI-based FC-matrices in calcium and hemodynamics both showed variability across time windows. On the other hand, FC matrices in calcium and hemodynamics within each time window were similar. If dFC in calcium and hemodynamics are matched, the similarity between calcium and hemodynamic FC in the same time window should be higher than that calculated using different time windows (e.g., similarity between Ca-FC_window#1_ and Hemo-FC_window#1_ would be higher than the similarity between Ca-FC_window#1_ and Hemo-FC_window#2_). Otherwise, the similarity between FC-matrices in calcium and hemodynamics merely reflects the overall similarity of FC in calcium and hemodynamics but not the coordinated “dynamics” of calcium and hemodynamic FC. Across all the data, we found that the distribution of the correlation coefficient between the FC-matrices in calcium and hemodynamics was shifted toward positive values compared with that calculated with the scan-shifted data (P < 10^-14^, Kolmogorov–Smirnov test; Fig. 2C). The difference between the real data and the trial-shifted data was also consistently positive across animals (P < 0.0156, n = 7 mice, sign rank test; Fig. 2D) and was seen across various window sizes ranging from 1 sec to 60 sec (Fig. 2E). Together these results suggest that temporal variability in hemodynamic FC, as measured with sliding window, arises from neural activity rather than from movement-related artifacts (Laumann et al. 2016) or non-neuronal physiological artifacts such as heartbeat and respiration (Bianciardi et al. 2009; Shmueli et al. 2007).

**Figure 2.**
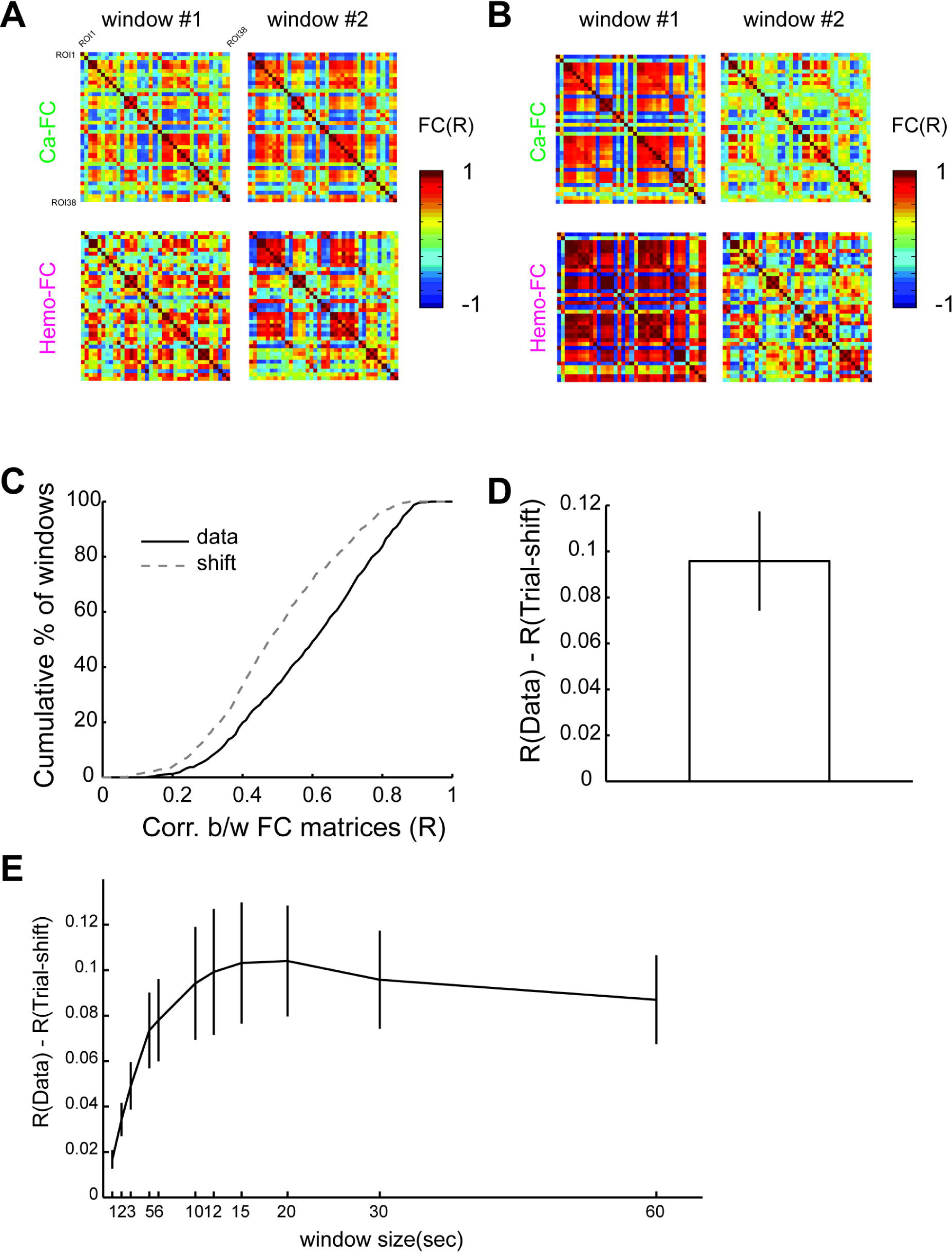
Significant relationship between calcium and hemodynamic FC calculated in short time windows. **(A)-(B)** Example ROI-by-ROI FC matrices for calcium and hemodynamics for different (non-overlapping) 30 sec windows. FC matrices were similar for calcium and hemodynamics in the same time window, but not across different time windows. (A) and (B) are two different examples from different animals. **(C)** Cumulative histogram of correlation between FC matrices for calcium and hemodynamics. Dotted line indicates trial-shifted control. **(D)** Correlation between FC matrices for calcium and hemodynamics was larger for the data than for the trial-shifted control significantly across animals. **(E)** Correlation between FC matrices of calcium and hemodynamics was larger for the data than the trial-shifted control across different window-sizes (1, 2, 3, 5, 6, 10, 12, 15, 20, 30 and 60 sec). Error bars indicate s.e.m. across animals (n = 7).

### Variations in transient neuronal coactivations explained variations in FC

What are the potential neuronal events that create dFC? Recent fMRI studies have proposed that variability in the neuronal coactivation pattern (CAP) of brain areas is directly reflected in the “dynamic” change of FC observed with the sliding window correlation (Liu and Duyn 2013). To address this possibility, for each scan, we compared the sliding window FC calculated using hemodynamics with the CAPs calculated using the calcium signal. The use of the calcium signal for extracting CAPs allowed us to capture faster spatiotemporal dynamics than the hemodynamics. More importantly, the use of two different signals also allowed us to avoid comparing sliding window FC and CAPs that were derived from the same signals and could lead to circular logic.

For each anatomical ROI, we first detected peaks in the calcium signal within a given time window and then defined CAPs as the frames in the calcium signal corresponding to the detected peak locations (Fig. 3A) (Liu and Duyn 2013). Similar to the previous reports in fMRI (Liu et al. 2013; Liu and Duyn 2013), we found variations in the spatial patterns of CAPs extracted from the same ROI (Fig. 3A, panels above time courses). We used CAPs to examine if whether variations of the spatial pattern of CAPs in different time windows could explain the spatial variation of sliding window FC. For each ROI in each 30 sec window, we extracted CAPs and FC using the calcium and hemodynamic signals, respectively. In the example 30 sec windows shown in Figure 3A, time courses of the chosen ROI showed transient activations that resulted in 11 and 3 frames of CAPs (corresponding to 1.1 and 0.3 sec of data, respectively). Despite the small number of frames corresponding to CAPs, the average spatial pattern of CAPs in the time window closely matched the spatial pattern of hemodynamic FC calculated in the same time window (compare the mean CAP and mean FC in Fig. 3A).

**Figure 3.**
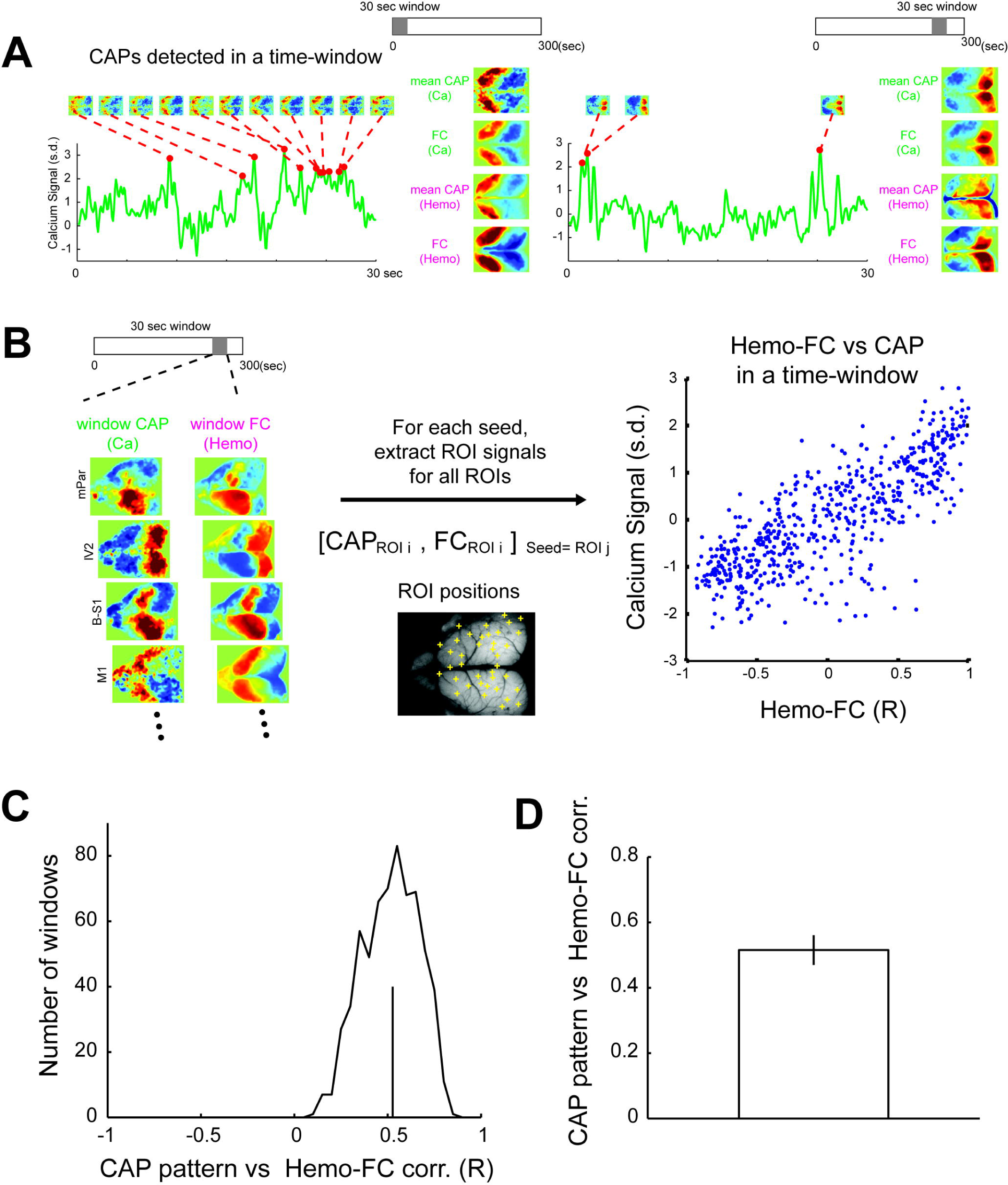
Comparison of calcium CAPs and hemodynamic FC across time-windows. **(A)** Procedure for detection of CAPs in calcium signal. For a given ROI, a calcium time course was extracted and z normalized (green time courses). Then, peaks exceeding 2 s.d. (red dots) were extracted. The frames corresponding to the peaks were considered CAPs (panels above the time courses). For each window, CAPs in calcium signal were averaged to obtain mean calcium CAP. Hemodynamic CAPs were calculated similarly (see Methods). Maps of Ca-FC and Hemo-FC were also calculated using the same time window. **(B)** Schematics to show the procedure of comparing calcium CAP and Hemo-FC across all ROI pairs in each time window. In each 30 sec time window, mean calcium CAPs and Hemo-FC maps were calculated for all ROIs as seeds (left). Then, for each seed-ROI j, calcium CAP and Hemo-FC values in ROI i were extracted to obtain a pair of CAP-FC values for the ROI-pairs (i, j) (middle). Finally, for each time-window, CAP-FC values were compared across all pairs of ROIs (right). **(C)** Histograms of correlation between CAP and Hemo-FC for all time windows across all animals. Vertical line indicates mean across time windows. **(D)** Mean correlation between CAP and Hemo-FC across animals. Note that (C) shows entire distribution of the data whereas (D) shows reproducibility across mice. Error bar indicates s.e.m. across animals (n = 7).

To further compare CAPs with sliding window hemodynamic FC across ROIs, we calculated CAPs for all pairs of ROIs and compared them with the FC of the same ROI-pairs in the same time window (Fig. 3B). Across all the data, CAP-matrices and FC matrices showed a high positive correlation (Fig. 3C–D; mean R = 0.525 across animals) suggesting that CAP and FC calculated using the same sliding window were similar.

The similarity between calcium CAP and hemodynamic FC in a short time window does not necessarily indicate coordinated temporal variation between CAP and FC, but could result entirely from similarity between the time-average patterns of CAP and FC. Therefore, to further examine whether coordinated temporal variations in CAPs and FC exist, we calculated ΔCAP and ΔFC by subtracting from each CAP and FC in each time window the average pattern of CAP and FC, respectively, that were calculated using the entire scan (Fig. 4A). A coordinated change in ΔCAP and ΔFC indicates a similar temporal fluctuation of CAP and sliding-window FC that cannot be accounted for by the similarity in the mean pattern of CAP and FC calculated using the entire scan. We found that the distribution of the correlation between ΔCAP and ΔFC for the real data was shifted toward positive values whereas the same distribution calculated using trial-shifted data was centered near zero (P < 10^-30^, Kolmogorov–Smirnov test; Fig. 4B). Furthermore, the correlation between ΔCAP and ΔFC was consistently positive across all animals (P < 0.156, n = 7 mice, sign rank test; Fig. 4C) and was seen across various sizes of time windows ranging from 1 to 60 sec (Fig. 4D). Excluding scans with potential artifacts using DVARS (Power et al. 2012; Power et al. 2014) still yielded a significant difference between the real and the trial-shifted data (Supplementary Fig. 9). Taken together, these results suggest temporal fluctuations of the spatial pattern of CAPs at least partly explain temporal fluctuations of hemodynamic FC calculated using sliding windows.

**Figure 4.**
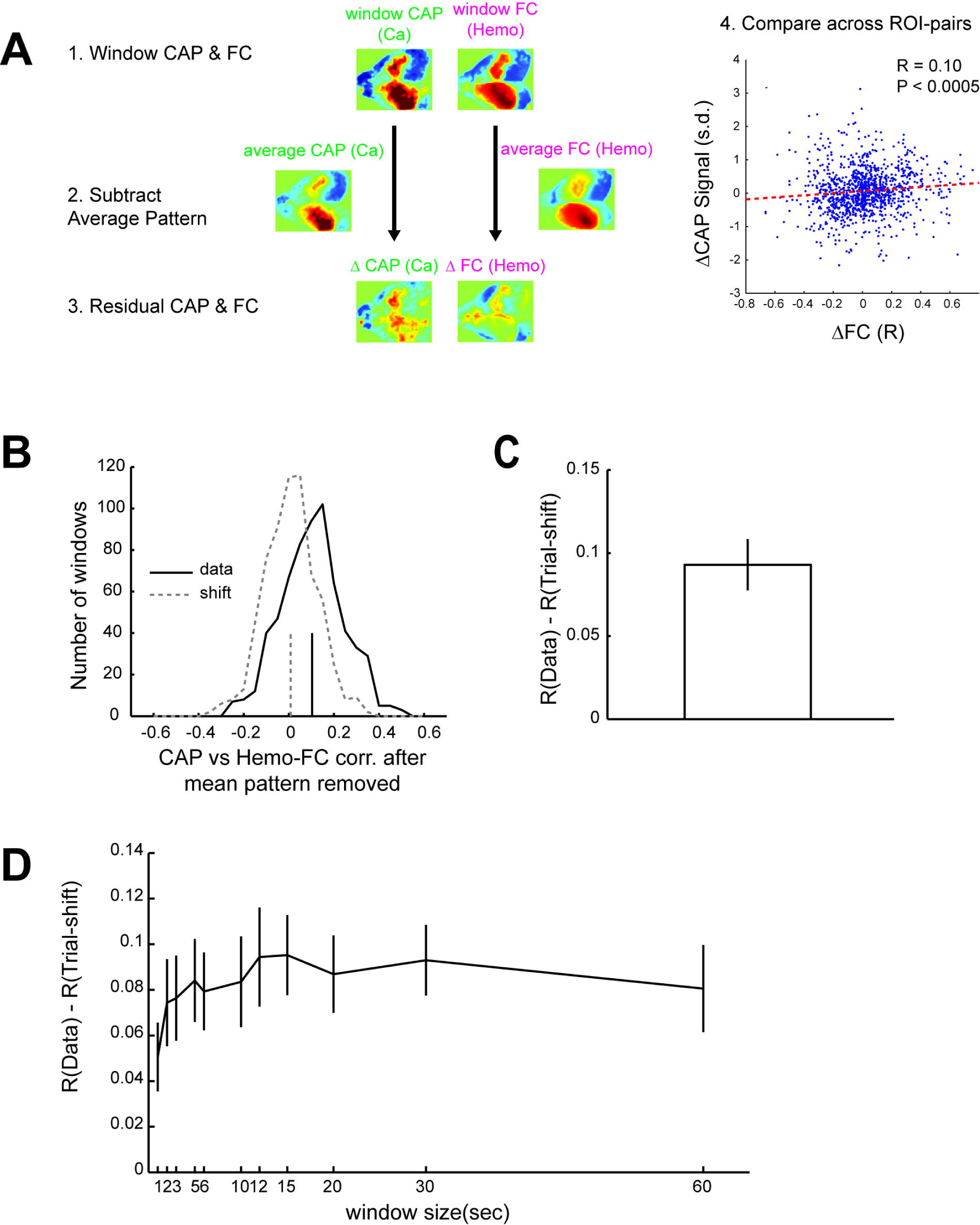
Temporal fluctuations in calcium CAPs and Hemo-FC was significantly related. **(A)** Schematics of the analysis. In each 30 sec time-window, mean calcium CAP and Hemo-FC were calculated (indicated as window CAP and window FC, respectively). From window CAP and window FC, average calcium CAP and average Hemo-FC that were calculated using the entire scan, in which the 30-sec window belongs to, were subtracted to obtain maps of ΔCAP and ΔFC, respectively. Finally, values of CAP and FC were compared across ROI pairs similarly as in Figure 3B. **(B)** Histograms of correlation between ΔCAP and ΔFC for all time windows across all animals. Vertical lines indicate mean across time windows. Solid and dotted lines indicate real and trial-shifted data, respectively. **(C)** Correlation between ΔCAP and ΔFC was significantly larger for the data than for trial-shifted control across animals. **(D)** Same as (C) but with different window-sizes (1, 2, 3, 5, 6, 10, 12, 15, 20, 30 and 60 sec). Error bars indicate s.e.m. across animals (n = 7).

### “Dynamics” of FC arise from non-stationarity of resting-state activity

Because FC is estimated by using a finite number of time-points, temporal fluctuations of FC observed in short time windows could arise from mere sampling error even when the underlying FC is stationary (Laumann et al. 2016). We next addressed whether the sampling error could explain the dFC observed in the present data. We compared two indices used in a previous study, namely cluster validity index and kurtosis, for real data and simulated data that were matched in spectral and covariance properties (Fig. 5A) (Laumann et al. 2016). The cluster validity index measures the degree of clustering of multiple sliding window FC calculated within the scan. Note that a smaller cluster validity index indicates more clustering (see Materials and Methods for details). For both calcium and hemodynamic signals, we found the cluster validity index of the real data to be significantly smaller than that of the simulated data (Fig. 5B), suggesting that the real data had a cluster structure that could not be fully accounted for by sampling error. Similarly, we calculated the kurtosis of the covariance matrices of real and simulated data. If the kurtosis of real data is larger than that of simulated data assuming a stationary Gaussian process, non-stationarity is implied for the real data, if the real data is normally distributed (Laumann et al. 2016). For the hemodynamic signal, the real data was approximately normally distributed (Supplementary Fig. 8A). Thus, we calculated the kurtosis using the hemodynamic signal without further preprocessing. For the calcium signal, because the original signal was not normally distributed, we applied further preprocessing before calculating the kurtosis. First, we applied an additional high pass temporal filter to remove low frequencies (< 0.1 Hz). Then we used a median filter to enforce the normality of the signal (Supplementary Fig. 6). For both hemodynamic and calcium signals, we found that the kurtosis of the real data was significantly higher than that of the simulated data (P < 10^-10^ for calcium, sign rank test, n = 50 scans that showed the approximately normal distribution of the signal (see Materials and Methods for the details of the assessment of normality); P < 10^-11^ for hemodynamics, sign rank test, n = 64 scans; Fig. 5C, Supplementary Fig. 8B). The calcium signal at low frequency (< 0.1 Hz) was approximately normal without additional preprocessing (Supplementary Fig. 7A). The kurtosis of the real and the simulated data was also significantly different at this frequency range, though the magnitude of the difference was smaller (Supplementary Fig. 7B). Together, these results suggest that dFC arises from the non-stationarity of spontaneous neuronal activity, and analyses based on sliding window correlation have the potential to detect non-stationarity.

**Figure 5.**
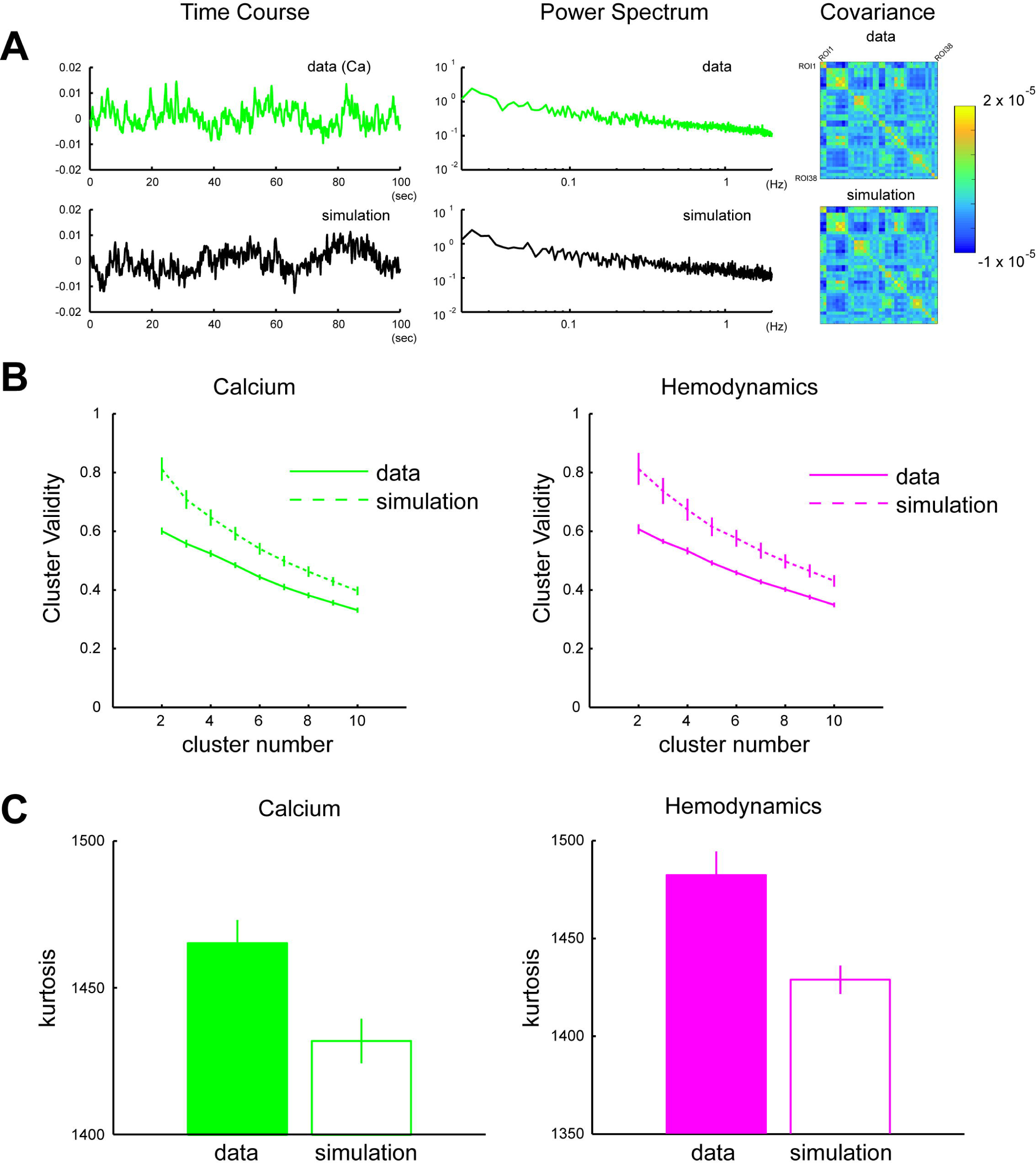
Comparison with simulated data indicated non-stationarity of the real data. **(A)** Examples of real and simulated time courses. Simulated time course (black) was matched to real data (green; calcium) in mean spectral content (middle panels) and ROI-by-ROI covariance matrix (right panels). The same procedure was applied to create simulated hemodynamic data (not shown). **(B)** Cluster validity index calculated for different number of clusters (k = 2-10). In both calcium (left) and hemodynamics (right), the cluster validity index was smaller for the real (solid lines) than the simulated data (dotted lines) indicating that the real data tended to be more clustered. **(C)** Kurtosis of real and simulated covariance matrices. For both calcium (left) and hemodynamics (right), multivariate kurtosis was larger for the real than for the simulated data. Error bars indicate s.e.m. across animals (n = 7).

To assess the potential contribution of motion-related and physiological artifacts on the analysis of the non-stationarity using kurtosis, we used DVARS calculated using the hemodynamic signal to exclude time points for which the artifacts might have been problematic (Power et al. 2012; Power et al. 2014). In some scans with large peaks in the time courses of DVARS, indicating potential body movements, the histogram of the DVARS was strongly right-skewed (Supplementary Fig. 3). Analysis of the kurtosis using a subset of scans for which the DVARS distribution was not right-skewed compared to the stationary simulation also revealed a significant difference between the real and the simulated data (P < 10^-14^, sign rank test, n = 25; Supplementary Fig. 4; see Materials and Methods for details). Furthermore, frame-censoring at several thresholds of DVARS revealed that the difference of kurtosis between the real and the simulated data did not depend on the levels of DVARS thresholds (Supplementary Fig. 5). These results suggest that the motion-related and physiological artifacts, as detected by DVARS, did not significantly affect the present results.

## Discussion

In the present study, we used simultaneous imaging of calcium and hemodynamic signals to show that temporal fluctuations in hemodynamic FC calculated in a short time window closely follow that of calcium FC, suggesting the neuronal origin of dFC. We have further shown that the spatial pattern of hemodynamic FC in a short time window is predicted by averaging the transient coactivations in the calcium signal (CAPs) contained within the same time window suggesting that temporally interspersed transient neuronal events underlie resting-state FC. Finally, we have shown that in both calcium and hemodynamic signals, statistical properties of FC calculated in a short time window were significantly different from those obtained with simulated signals that were stationary by construction. These results advocate for the analysis of the “dynamic” aspect of FC obtained in human fMRI experiments. Insights of the neuronal events underlying dFC provided by the present study would also be informative for developing appropriate analysis methods for dFC.

### Relationship to previous investigations of the neuronal origin of dFC

To provide direct evidence linking neuronal activity and dFC, several groups have conducted simultaneous recording of fMRI and local field potential (LFP) (Lu et al. 2007; Pan et al. 2011; Thompson et al. 2013) or EEG (Chang et al. 2013; Tagliazucchi et al. 2012b). However, these previous studies were limited in several ways. Because LFP recordings were limited by a small number of recording sites and EEG recordings did not have enough spatial resolution, evidence directly linking the global spatial pattern of neuronal activity with hemodynamic FC has been lacking. Using simultaneous imaging of calcium and hemodynamic signals, the present study provides evidence suggesting that temporal variability of hemodynamic FC and its time-to-time spatial patterns reflect spatial patterns of large-scale neuronal activity. Moreover, because the present study used anesthetized and head-fixed mice, the results are unlikely to be attributable to head motion.

Recent human fMRI studies have proposed that neuronal activity important for FC is condensed into transient large scale neuronal coactivations (i.e. CAPs) (Liu and Duyn 2013; Tagliazucchi et al. 2012a; Tagliazucchi et al. 2011). Consistent with this idea, imaging studies in mice have revealed transient neuronal coactivations across brain areas (Matsui et al. 2016; Mohajerani et al. 2013; Vanni and Murphy 2014). In our previous study, we searched for neuronal coactivations that resembled the spatial patterns of (static) FC and showed that such neuronal coactivations were converted into spatially similar hemodynamic signals (Matsui et al. 2016). In the present study, we took a different approach that was similar to single frame analysis methods employed in recent human fMRI studies (Karahanoğlu and Van De Ville 2015; Liu et al. 2013; Liu and 2013; Tagliazucchi et al. 2011). We were especially interested whether CAPs represent the potential neuronal events underlying the temporal variability of sliding window FC. Because derivation of CAPs and sliding-window FC using identical BOLD signals potentially lead to circular logic, in the present study, we examined the link between CAPs and sliding window FC derived from different signals (calcium and respectively). Instead of specifically looking at neuronal coactivations that resembled “static” FC, we included all the individual CAPs into our analysis and showed that variation of the spatial pattern of individual CAPs across time windows was significantly related to variations of hemodynamic FC across time windows. Thus, the present findings suggest the importance of the development of an analysis that specifically focuses on CAPs (Karahanoğlu and Van De Ville 2015; Liu et al. 2013). However, the modest correlation found between CAP and sliding window FC implies that the fluctuations of calcium and hemodynamic signals from the average pattern maybe small. It should also be noted that, although statistically significant, the correlation between ΔCAP and ΔFC was relatively weak. Part of the reason for this could be non-neuronal physiological noise that contributed to the hemodynamics (Matsui et al. 2016). In the present study, because of the use of anesthesia and head-fixation, head motion was is unlikely to be the primary source of the non-neuronal noise. However, other physiological activities (e.g. respiration and heartbeat) are known to affect hemodynamics (Chang et al. 2009; Chang and Glover 2009) and, thus, are likely to affect temporal fluctuation of hemodynamic FC as well. Although scan-wise data exclusion using DVARS suggested that the motion-contamination was not the major cause of the observed temporal correlation between ΔCAP and ΔFC (Supplementary Fig. 9), the common artifacts on hemodynamic and calcium signals could have contributed to the observed temporal correlation. Our results (i.e. relatively low correlation between ΔCAP and ΔFC) indicate that correction for such non-neuronal physiological noise (Glover et al. 2000) is likely to be essential for the analysis of dFC.

### Non-stationarity of spontaneous brain activity correlation

It has been a matter of debate as to what extent temporal fluctuations of FC are attributed to the dynamics of underlying neuronal activity and not to non-neuronal sources of noise (e.g., head motion, sampling variability; reviewed in (Hutchison et al. 2013)). Laumann and colleagues have reported that most of the temporal fluctuations of single subject FC is explained by head motion (Laumann et al. 2016). After controlling for the head motion, Laumann and colleagues have concluded that the statistical properties of resting-state FC in human fMRI is are indistinguishable from those obtained with simulated signals that are stationary by construction. A similar study by Hindriks and colleagues has also indicated that the apparent dFC calculated with the sliding window method does not necessarily indicate non-stationarity of the resting brain network (Hindriks et al. 2016). However, in terms of spontaneous neuronal activity itself, there is substantial evidences showing that spontaneous neuronal activity is non-stationary (Foster and Wilson 2006; Ji and Wilson 2007; Logothetis et al. 2012). In particular, under both awake and anesthetized states, transient neuronal events such as sharp wave ripples have been shown to produce coordinated activity across the entire brain (Logothetis et al. 2012). The present results are consistent with these previous studies supporting the non-stationarity of neuronal activity and further showed that FC calculated using such non-stationary neuronal activity also showed non-stationarity, as expected.

### Potential contribution of arousal

Fluctuation in the level of arousal has been shown to contribute to the apparent “dynamics” and non-stationarity of FC (Laumann et al. 2016; Tagliazucchi and Laufs 2014). The present study observed larger temporal variability of hemodynamic FC in anesthetized mice than in awake, eyes open-fixated humans (Laumann et al. 2016). Interestingly, a recent study reported large within-subject FC variability in subjects instructed to close their eyes (Pannunzi et al. 2017), which predisposes them to sleep (Tagliazucchi and Laufs 2014). In the present study, it could be possible that a fluctuating level of anesthesia, instead of the subject’s sleep state, could have contributed to the greater variability of FC. Alternatively, rapid fluctuation between awake and sleep states in mice (Adler et al. 2014) could have contributed to the greater variability of FC compared to humans. According to the results of the analysis using power spectra (Supplementary Fig. 2) and DVARS (Supplementary Figs. 3–5), we consider that the level of arousal rarely reached a point at which the animal started to struggle. Nevertheless, the present data alone were not sufficient to exclude the possibility that the fluctuation of the level of arousal accounted for a large fraction of the FC “dynamics” and non-stationarity observed here. Future experiments conducting simultaneous recordings of functional images and physiological signals (e.g. electroencephalogram, respiration-rates) in mice would be able to assess the exact amount of non-stationarity in FC under a defined state of arousal.

### Other limitations of the study

It should be clearly stated that the present results do not guarantee that sliding window methods are always capable of detecting non-stationarity in human resting-state fMRI data. The present study used tightly head-restrained animals and high signal-to-noise-ratio (SNR) imaging at a high frame rate (5 and 10 Hz for the hemodynamics and calcium signal, respectively). Compared to the present experimental conditions, overall SNR in typical human resting-state fMRI is likely to be substantially compromised. Under such low SNR conditions, it is not clear whether simple sliding window correlation methods can detect the non-stationarity of FC (Hindriks et al. 2016; Laumann et al. 2016). With respect to SNR, we expect that the recent development of high-speed fMRI (Feinberg et al. 2010) will significantly improve the detectability of non-stationarity. Nevertheless, the present results suggest that, rather than the sliding window based method, an alternative analysis strategy that directly extracts CAPs from hemodynamic signals (Karahanoğlu and Van De Ville 2015; Liu et al. 2013; Liu and Duyn 2013; Tagliazucchi et al. 2012a) may be more appropriate for extracting relevant information related to dFC. Care should be taken, however, because the smaller brain and the low dimensionality of FC in the mouse (compared to the human subjects) could also have made CAPs from just a few frames look similar to those of FC.

It should also be noted that the present results do not claim that dFC has significant behavioral or cognitive consequences. Instead of examining the potential relationship between dFC and cognitive dynamics or behavioral variability [see (Preti et al. 2016) for a recent review], here we focused on validating the neuronal origin of dFC. Experiments under anesthesia greatly reduced potential confounding factors, such as head motion and arousal state (Hutchison et al. 2014; Laumann et al. 2016).

Nevertheless, the present wide-field imaging setup can be naturally extended to awake imaging with task-performing mice (Ferezou et al. 2007; Wekselblatt et al. 2016). Such experiments would reveal the potential consequences of dFC on the behavior.

## Acknowledgements

We thank A. Honda and Y. Sono for animal care and genotyping; the Research Support Center, Graduate School of Medical Sciences, Kyushu University, for technical support: and K. Jimura and Y. Noro for helpful discussion. Support for this work was provided by grants from Brain Mapping by Integrated Neurotechnologies for Disease Studies (Brain/MINDS) – Japan Agency for Medical Research and Development (AMED), Core Research for Evolutionary Science and Technology (CREST) – AMED and Strategic International Research Cooperative Program (SCIP) – AMED, and Japan Society for Promotions of Sciences (JSPS) KAKENHI Grants 25221001 and 25117004 (to K.O.), World Premium Institute (WPI), JSPS (to K.O.); JSPS KAKENHI Grant 17K14931 (to T. Matsui) and JSPS Research Fellowship 20153597 (to T. Murakami).

## Figure Captions

**Supplementary Figure 1.**
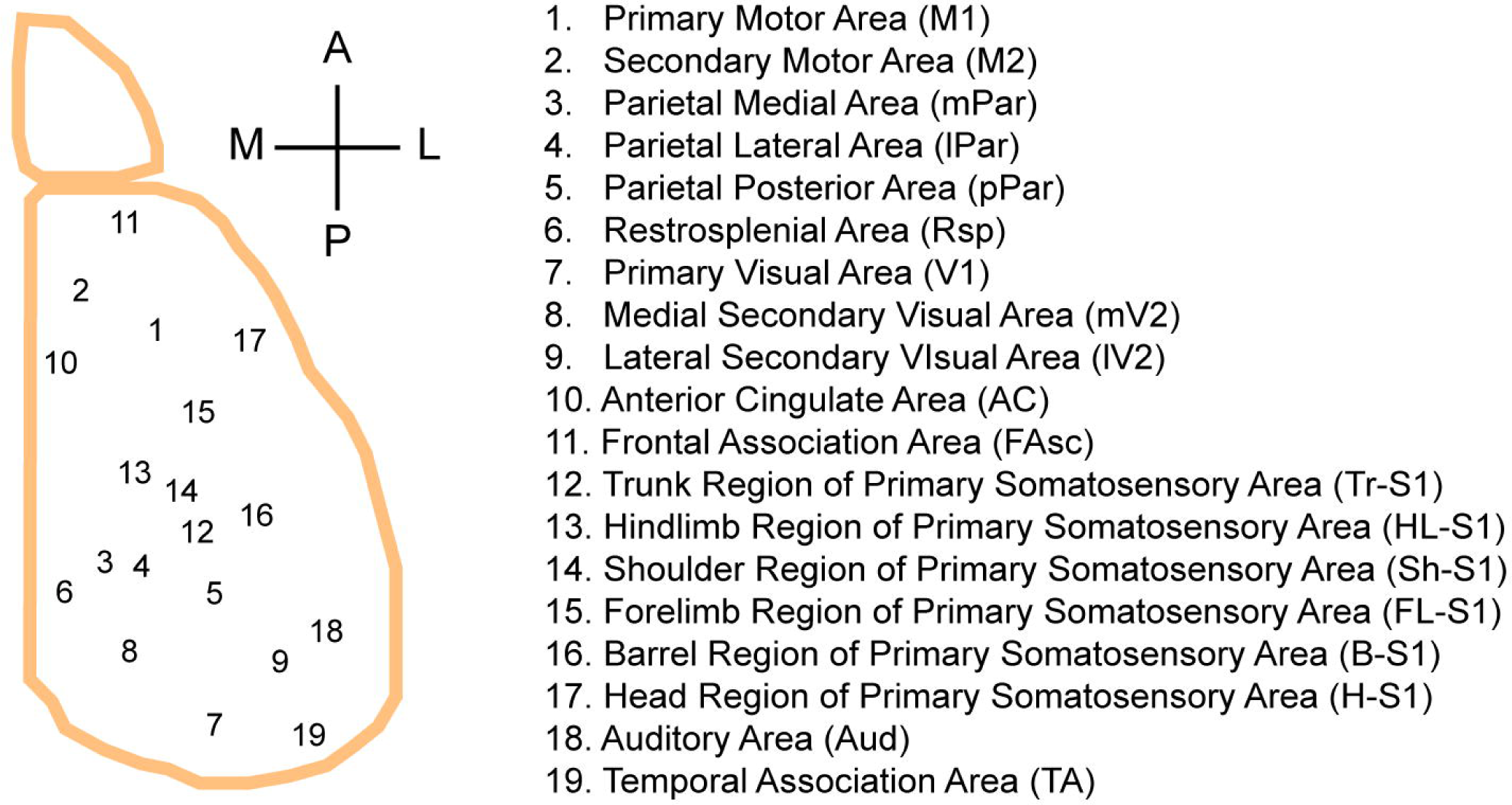
Anatomical locations of ROIs. Anatomical locations of 19 ROIs are shown for the right hemisphere. Anatomical nomenclatures of ROIs are shown on the right. ROIs in the left hemisphere are taken at mirror symmetric positions to yield a total of 38 ROIs.

**Supplementary Figure 2.**
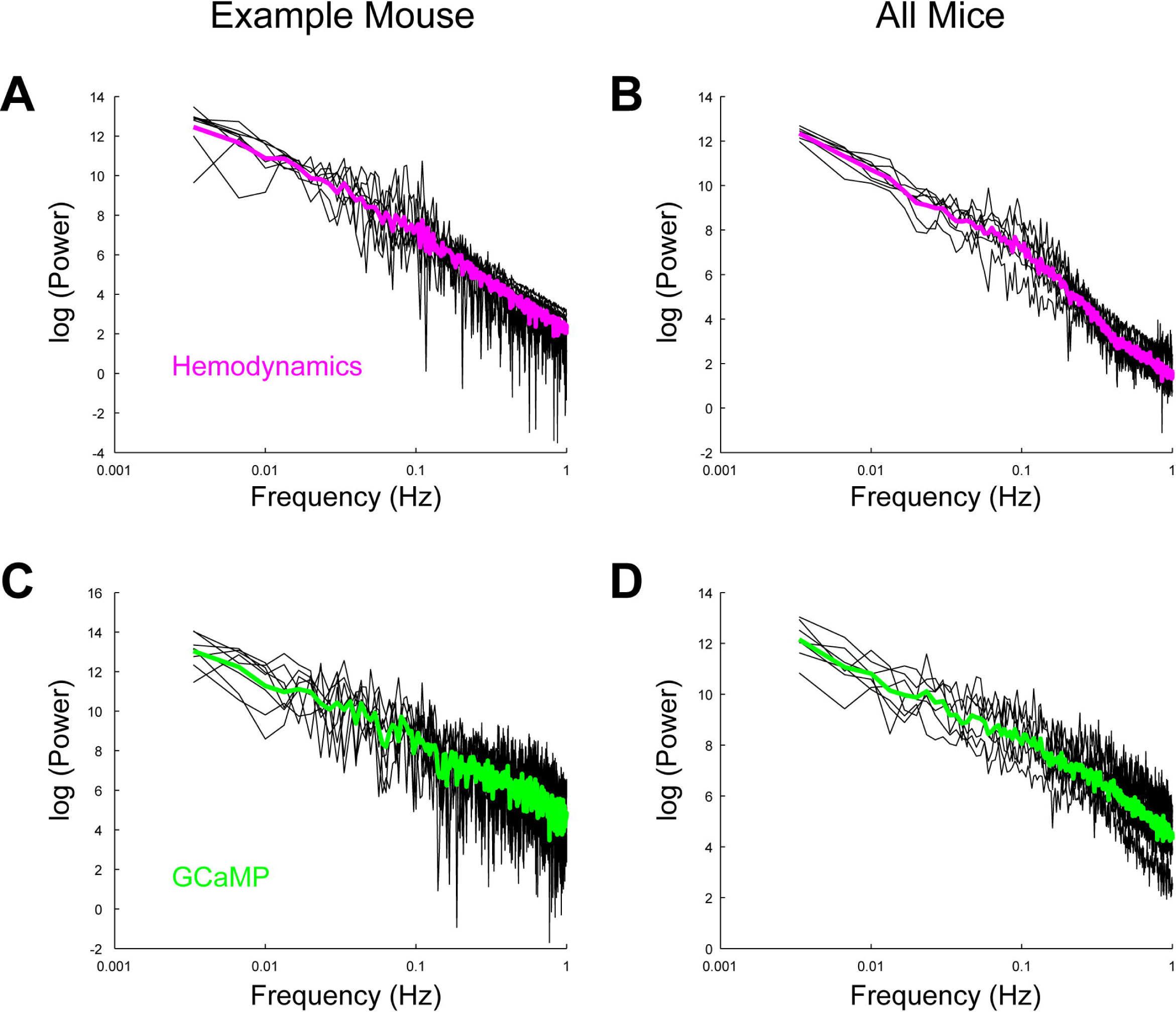
Power spectrum of calcium and hemodynamic signals. **(A)** Example power spectrum of hemodynamic signal for one mouse. Hemodynamic signals were not temporally filtered. Black, individual scan. Red, average across scans. **(B)** Average power spectrum of hemodynamic signal across mice (Red). Black, individual mouse (averaged across scans). **(C)** Example power spectrum of calcium signal for one mouse. Calcium signals were not temporally filtered. Black, individual scan. Red, average across scans. **(D)** Average power spectrum of calcium signal across mice (red). Black, individual mouse (average across scans).

**Supplementary Figure 3.**
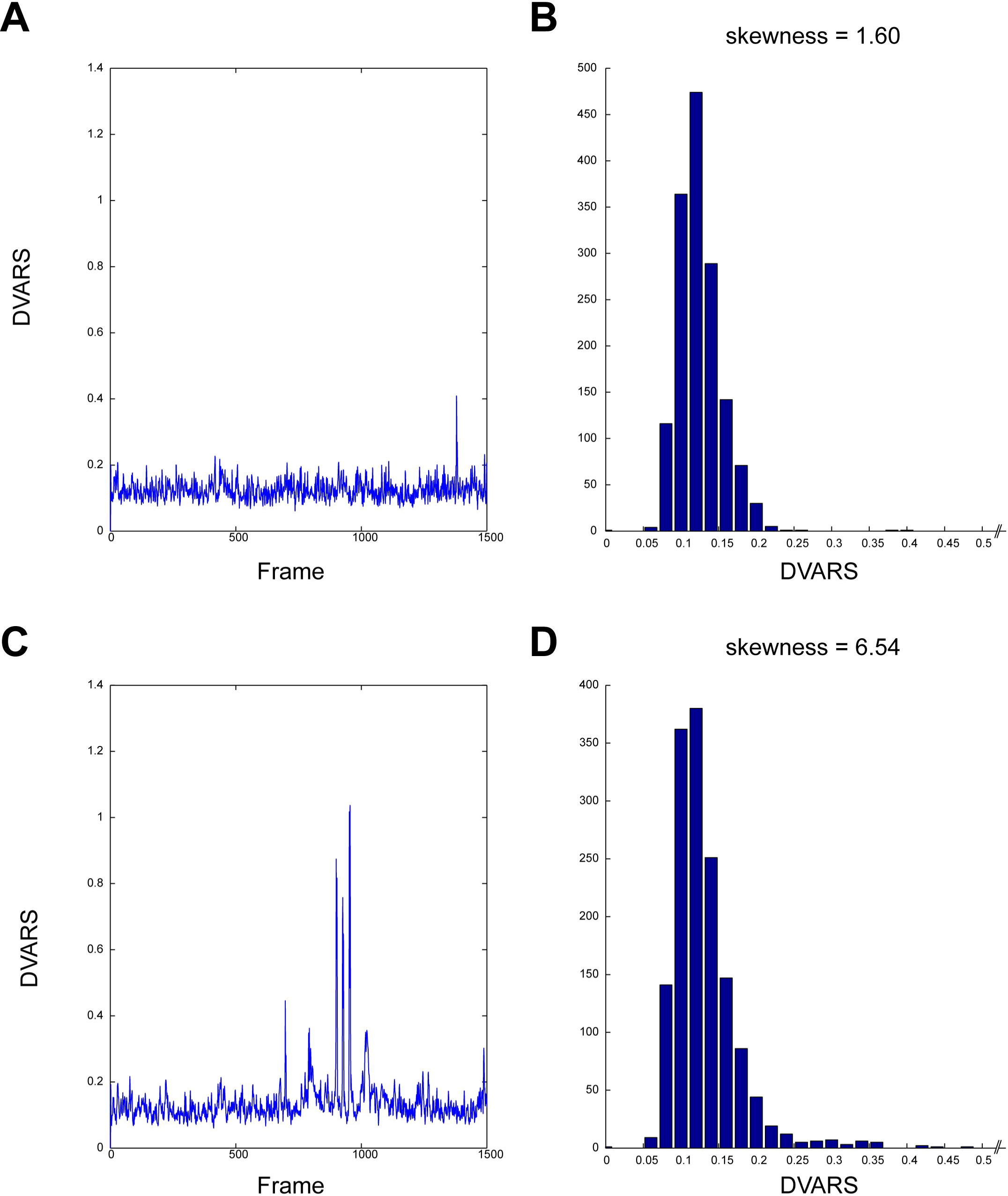
Example time courses and histograms of DVARS calculated using hemodynamic signal. **(A)** Example time course of DVARS for one mouse. **(B)** Histogram of DVARS for the data shown in (A). Inset, same histogram but for different y-axis range. The histogram was not markedly right skewed (skewness = 1.60). **(C)** Another example time course of DVARS obtained in the same mouse as in (A). In this scan, large peaks probably representing motion and/or physiological artifacts were observed. **(D)** Histogram of DVARS calculated using data shown in (C). Inset, same histogram but for different y-axis range. The histogram was strongly right-skewed (skewness = 6.54).

**Supplementary Figure 4.**
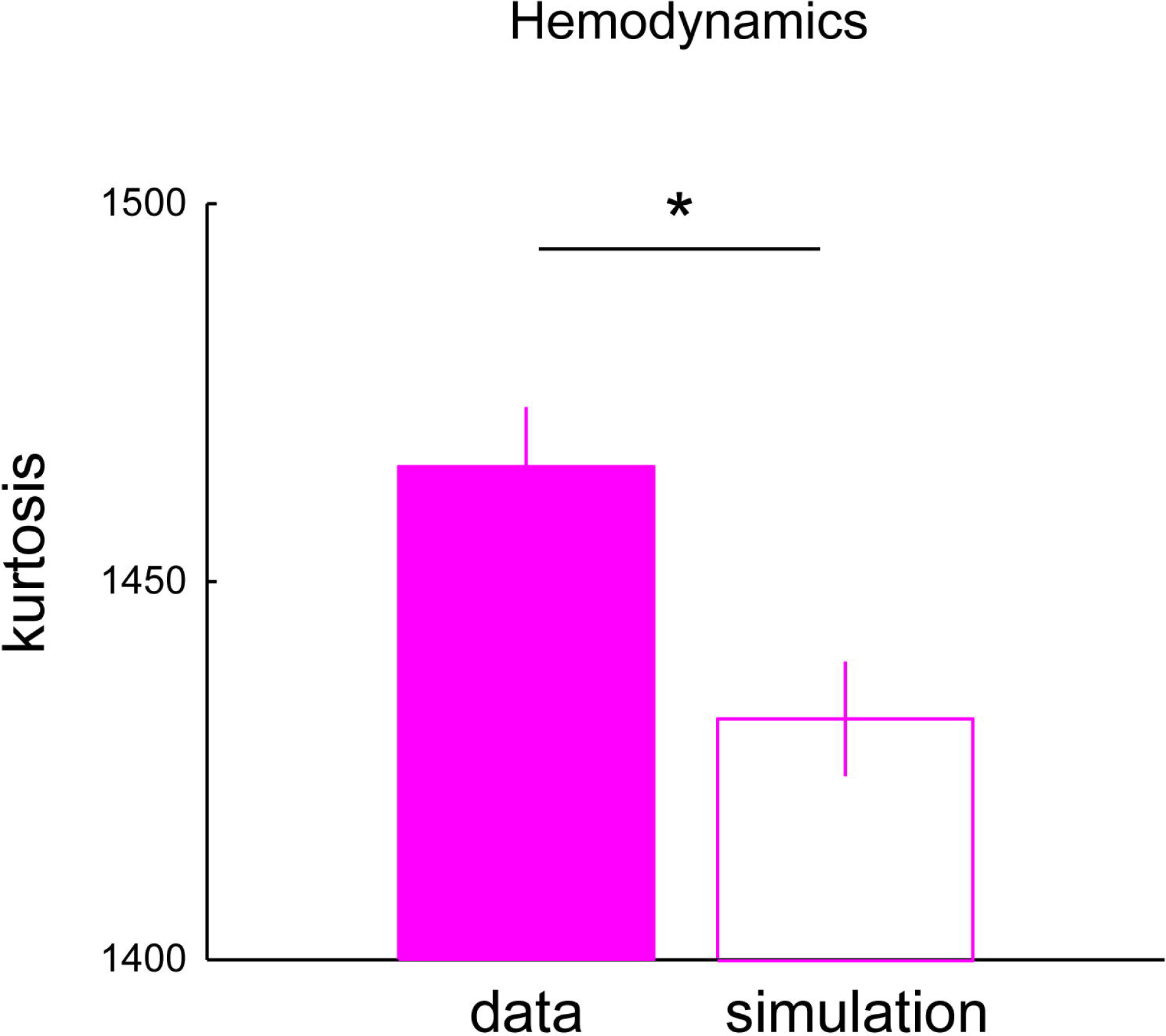
Difference between the kurtosis of real and simulated data in Hemodynamics. Kurtosis was calculated for scans selected based on the distribution of DVARS (n = 25). See Methods for details of the selection of the scans. *, P < 0.0001 (sign rank test). Error bars indicate s.e.m.

**Supplementary Figure 5.**
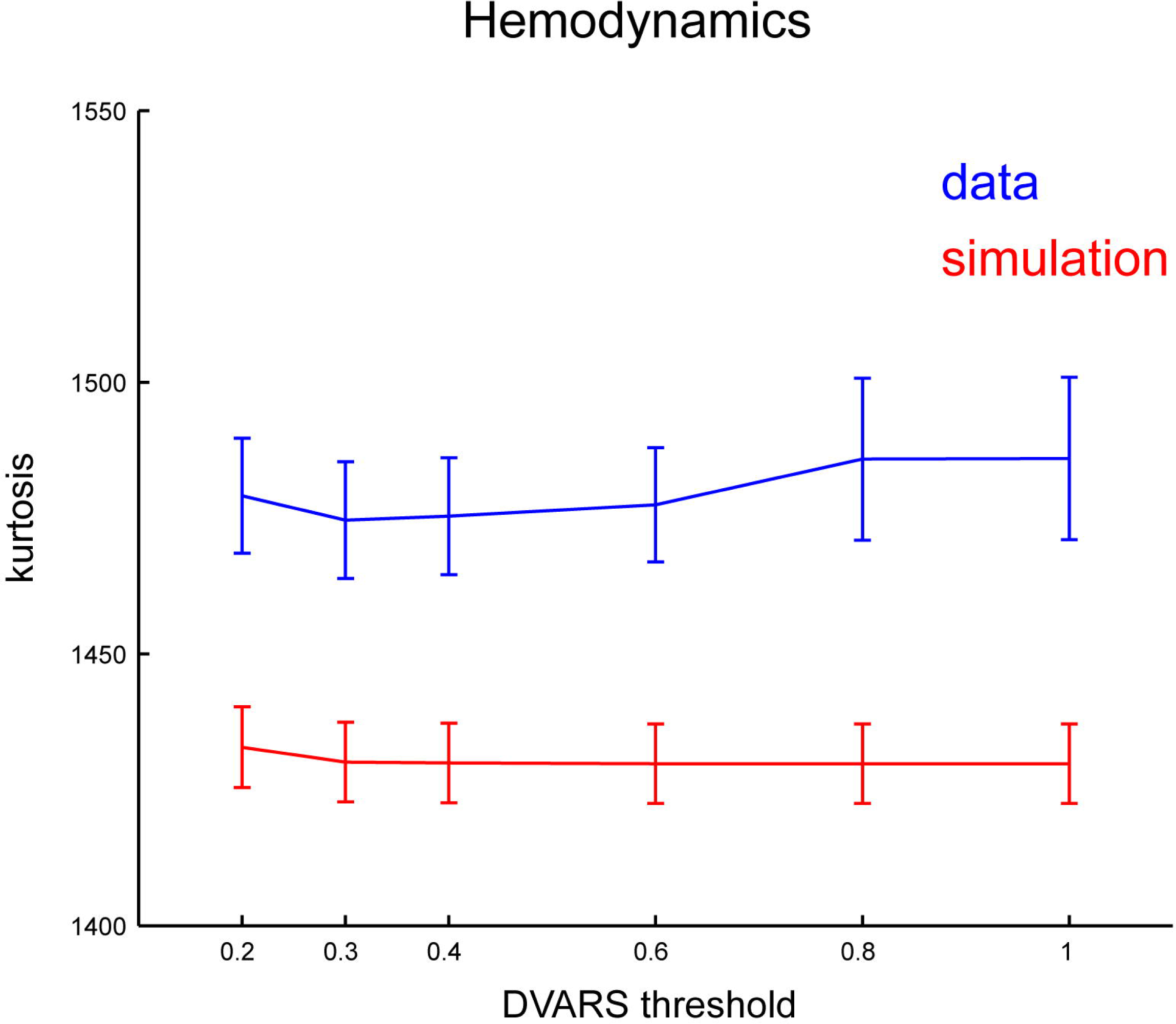
Difference between the kurtosis of real and simulated data in hemodynamics as a function of DVARS threshold for frame-censoring. The difference of the kurtosis between the real and the simulated data (blue and red, respectively) was largely insensitive to the level of the DVARS threshold. Note that the smaller DVARS threshold indicates stricter threshold. Error bars indicate s.e.m.

**Supplementary Figure 6.**
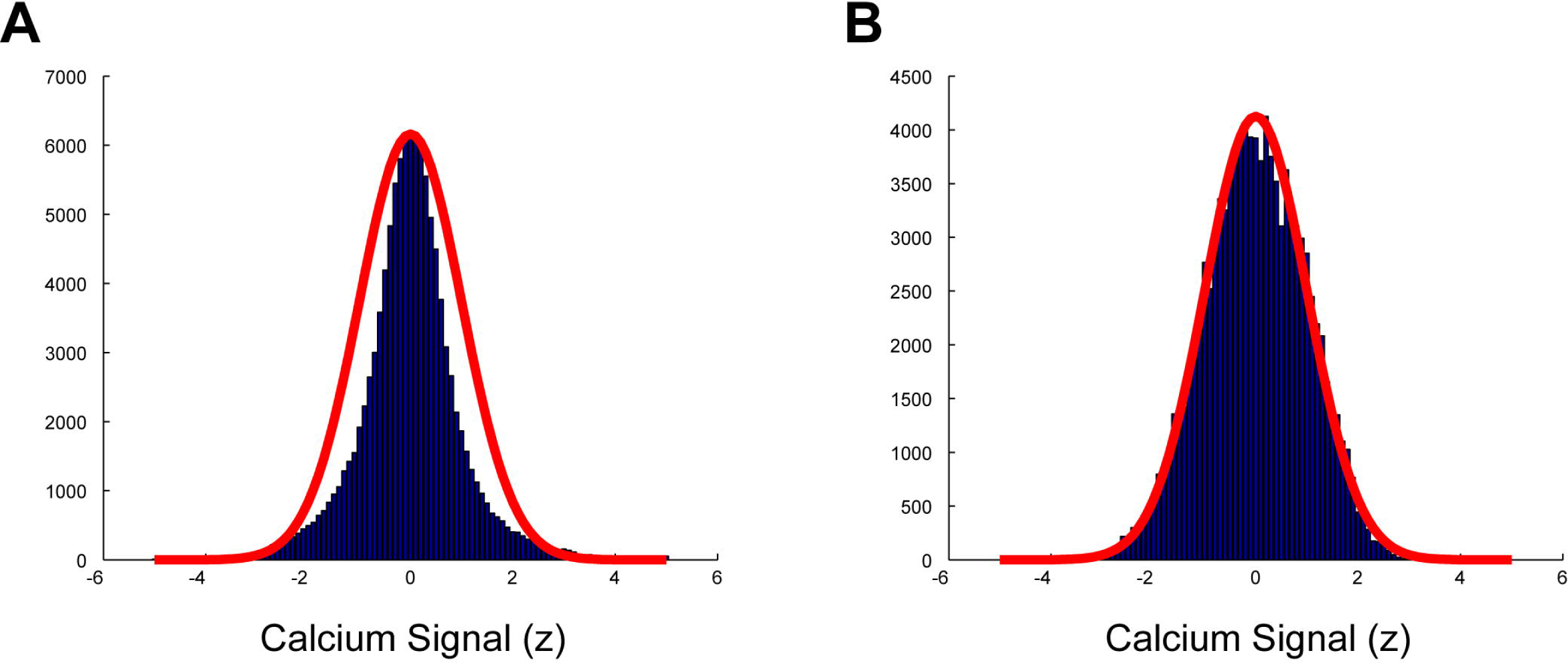
Kurtosis analysis using calcium signal processed with additional high-pass and media filters. **(A)** Example histogram of high frequency calcium signal (> 0.1 Hz) in one ROI of one mouse before the application of median filter. Red, Gaussian fit. **(B)** Same as (A) but for the signal after the application of median filter. Red, Gaussian fit. Note better fitting with Gaussian after median filter application, suggesting enforced normality of the signal.

**Supplementary Figure 7.**
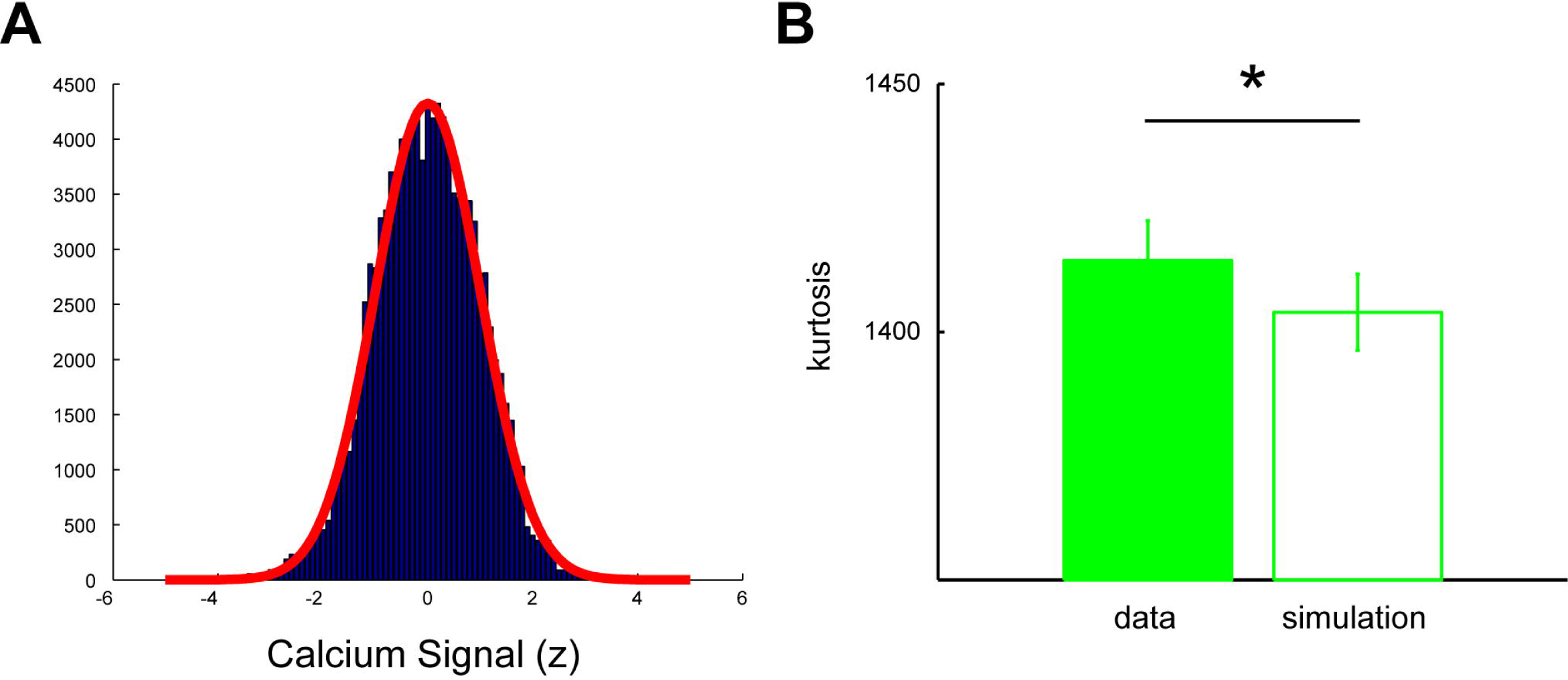
Kurtosis analysis using low frequency calcium signal. **(A)** Example histogram of low frequency calcium signal (0.01 Hz < f < 0.1 Hz) in one ROI of one mouse. Red, Gaussian fit. (B) Difference between the kurtosis of real and simulated data in low frequency calcium signal for scans selected based on the normality of the signal (55 scans). *, P < 10^-9^, sign rank test. Error bars indicate s.e.m.

**Supplementary Figure 8.**
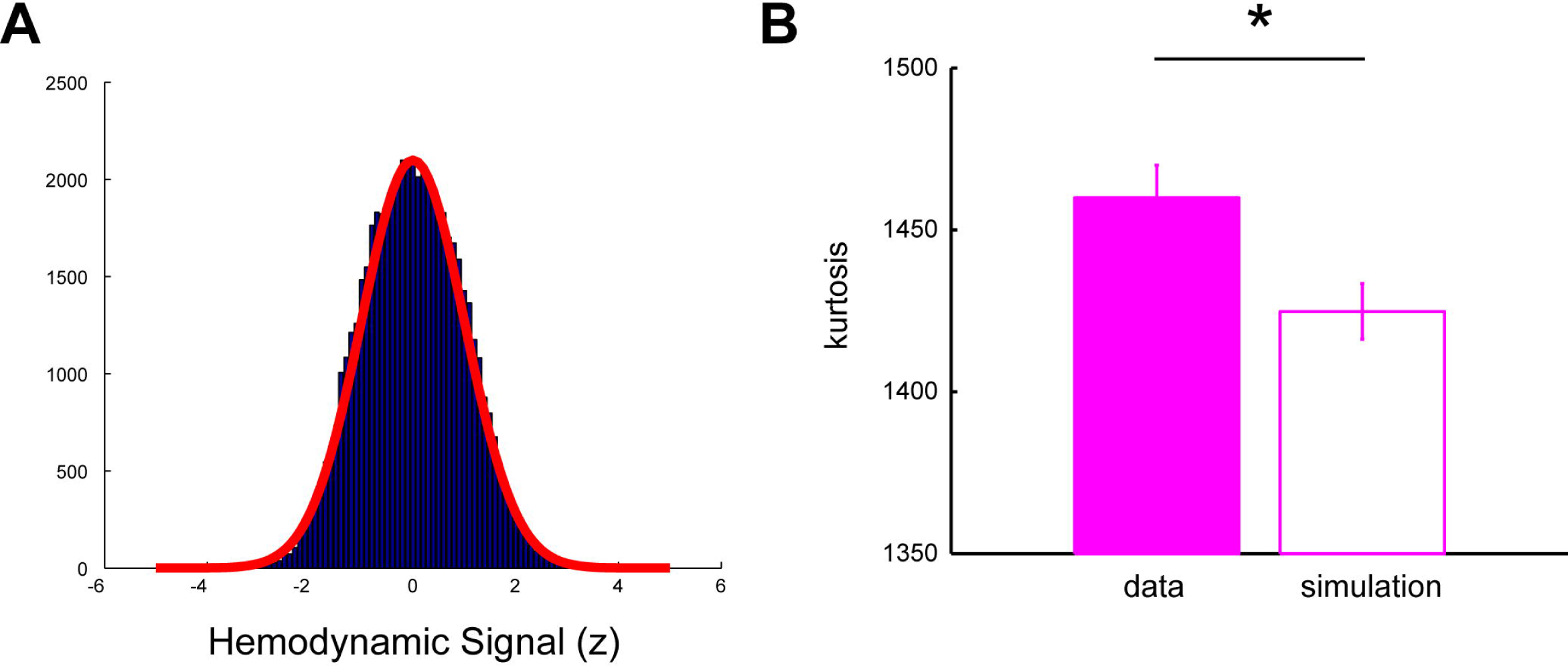
Hemodynamic signal was approximately normally distributed. **(A)** Example histogram of hemodynamic signal in one ROI of one mouse. Red, Gaussian fit. Note that no additional preprocessing was performed. **(B)** Difference between the kurtosis of real and simulated data in hemodynamic signal for scans selected based on the normality of the signal (45 scans). *, P < 10^-8^, sign rank test. Error bars indicate s.e.m.

**Supplementary Figure 9.**
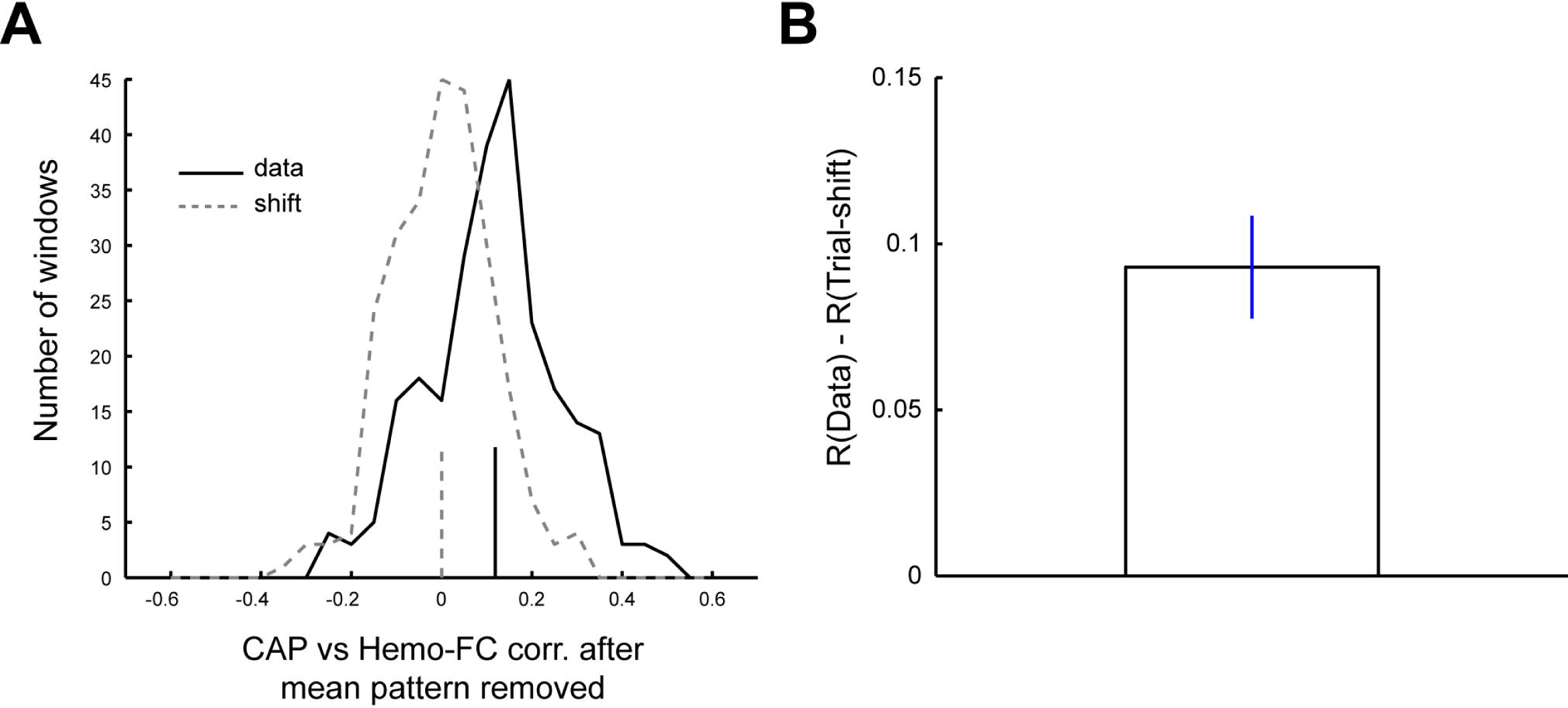
Significant temporal correlation between CAP and windowed-FC remained after removing scans with potential artifacts. **(A)** Same convention as in Figure 4B but for the data using scans selected for small movement and/or physiological artifacts based on DVARS (25 scans). *, P < 10^-17^, Kolmogorov-Smirnov test. **(B)** Same convention as in Figure 4C but for the selected scans. Positive difference between the real and the shifted data was observed in 5 out of 7 mice.

